# A brainstem to hypothalamic arcuate nucleus GABAergic circuit drives feeding

**DOI:** 10.1101/2023.05.03.538908

**Authors:** Pablo B Martinez de Morentin, J Antonio Gonzalez, Georgina K.C. Dowsett, Yuliia Martynova, Giles S.H. Yeo, Sergiy Sylantyev, Lora K Heisler

## Abstract

The obesity epidemic is principally driven by the consumption of more calories than the body requires. It is therefore essential that the mechanisms underpinning feeding behavior are defined. The brainstem nucleus of the solitary tract (NTS) receives direct information from the digestive system and projects to second order regions in the brain. Though γ-Aminobutyric acid is widely expressed in the NTS (GABA^NTS^), its function has not been defined. Characterization of GABA cells using single nucleus RNA sequencing (Nuc-Seq) identified at least 19 clusters. Here we provide insight into the function of GABA^NTS^ cells, revealing that selective activation of GABA^NTS^ neurons significantly controls food intake and body weight. Optogenetic interrogation of GABA^NTS^ circuitry identified GABA^NTS^→arcuate nucleus of the hypothalamus (ARC) projections as appetite suppressive without creating aversion. Electrophysiological analysis revealed GABA^NTS^→ARC stimulation inhibits hunger promoting agouti-related protein/neuropeptide Y (AgRP/NPY) neurons via GABA release. Adopting an intersectional genetics strategy, we clarify that the GABA^NTS^→ARC circuit induces satiety. These data identify GABA^NTS^ as a new modulator of feeding behavior, body weight and controller of orexigenic AgRP/NPY activity, thereby providing insight into the neural underpinnings of obesity.

**Highlights:**

- Nucleus of the solitary tract (NTS) GABA neurons are responsive to nutritional status.
- Chemogenetic GABA^NTS^ neuron activation reduces food intake and body weight.
- GABA^NTS^ projections to the hypothalamic arcuate nucleus (ARC) promote satiety.
- Optogenetic GABA^NTS^→ARC stimulation inhibits orexigenic AgRP/NPY neurons.

**In Brief:** Martinez de Morentin et al. identify GABAergic neurons in the nucleus of the solitary tract as a new player in the circuit governing feeding behavior and body weight.

## Introduction

Obesity represents a key challenge to human health and is primarily due to the consumption of calories in excess to body’s energy requirements. Eating is a complex behavior that not only depends on the basic energy demands at a cellular and organism level, but also the integration of internal and environmental cues, the reward value of food, motivation, and conditioning behavior (Andermann & Lowell, 2017; Campos et al., 2022). The aim of the present study was to probe neurocircuitry regulating feeding and body weight with the objective of uncovering critical energy homeostasis circuitry.

One of the primary nodes for the integration of energy-related information from the periphery to the brain is the nucleus of the solitary tract (NTS) within the brainstem dorsal vagal complex (DVC)(Hyun & Sohn, 2022). The NTS contains a heterogeneous population of energy-related sensitive cells(Cheng et al., 2022; Dowsett et al., 2021; Grill & Hayes, 2012). Recent studies using single-nucleus RNA sequencing provide a detailed expression map of identified cellular populations involved in energy balance, and most were revealed to be glutamatergic(Dowsett et al., 2021; Ludwig et al., 2021). For example, subpopulations of these glutamatergic cells influencing energy balance include leptin receptor (LEPR)(Cheng et al., 2020), calcitonin receptor (CALCR)(J. Chen et al., 2020a), glucagon-like peptide 1 receptor (GLP-1R)(Alhadeff et al., 2017; Fortin et al., 2020), preproglucagon (PPG)(Holt et al., 2019), tyrosine hydroxylase (TH) (Aklan et al., 2020; J. Chen et al., 2020b), cholecystokinin (CCK) (D’Agostino et al., 2016) and proopiomelanocortin (POMC) (Georgescu et al., 2020; Zhan et al., 2013). However, very little is known about the role of NTS inhibitory GABA-releasing clusters (GABA^NTS^) in energy homeostasis. One report in rats indicated a subpopulation of GLP-1R-expressing neurons releasing GABA are necessary mediators for the anorectic effects of the obesity medication liraglutide (Fortin et al., 2020). Therefore, GABA^NTS^ represents and intriguing and understudied population of NTS cells and is the focus of the present study.

In the regulation of feeding behavior, one of the most widely studied projections from the NTS is to the hypothalamus (Aklan et al., 2020; Blevins et al., 2004; D’Agostino et al., 2016; Liu et al., 2017; Shi et al., 2021; Tsang et al., 2020). However, whether the arcuate nucleus of the hypothalamus (ARC) receives GABAergic inhibitory control from the NTS is not known. Within the ARC is a subpopulation of potent orexigenic neurons expressing both agouti-related peptide and neuropeptide Y (AgRP/NPY) (Betley et al., 2013; Hahn et al., 1998;

Heisler et al., 2006; Heisler & Lam, 2017). We hypothesized that GABA^NTS^ neurons significantly regulate feeding and body weight and project to and inhibit key hunger-stimulating AgRP/NPY cells.

## Results

### GABA^NTS^ cells are sensitive to energy status and modulate food intake

To characterize GABA^NTS^ neurons, we initially used a single nucleus RNA sequencing (NucSeq) dataset of the mouse dorsal vagal complex (Dowsett et al., 2021). Neuronal cells expressing transcripts for *Slc32a1* (solute carrier family 32 member 1 or vesicular GABA transporter, Vgat) were extracted and re-clustered, consisting of 1847 neurons, which formed 19 clusters (**Figure 1A**). Of these clusters, 2-Cacna2d1/Tmem163 expressed Vglut2 transcripts, and lower levels of *Gad1* and *Gad2.* Low expression of classical neurotransmitter related genes was found in these GABAergic neuronal clusters (**Figure 1B**). Adipocyte hormone Leptin receptor (*Lepr)* expression was identified in 13-Onecut2/Lepr and 15-Ebf2/Acly clusters (**Figure 1B**). We identified some expression of incretin receptors including glucagon-like receptor 1 (GLP-1R) and gastric inhibitory polypeptide receptor (GIPR) in several clusters (**Figure S1A**). Differential gene expression analysis was performed on each cluster to identify the effects of an overnight fast on transcript expression in GABAergic (*Slc32a1*+) neurons (**Figure S1B**). Cluster 3-Dcc/Cdh8 displayed significant upregulation in transcript expression in response to a fast, however 93.7% of this cluster originates from *ad libitum* fed animals. Significantly differentially regulated genes in *Slc32a1^+^* neurons can be found in **Figure S1C**. These data indicate minimal overlap with specific populations of NTS neurons that have been previously described with regards to energy balance.

**Figure 1.**
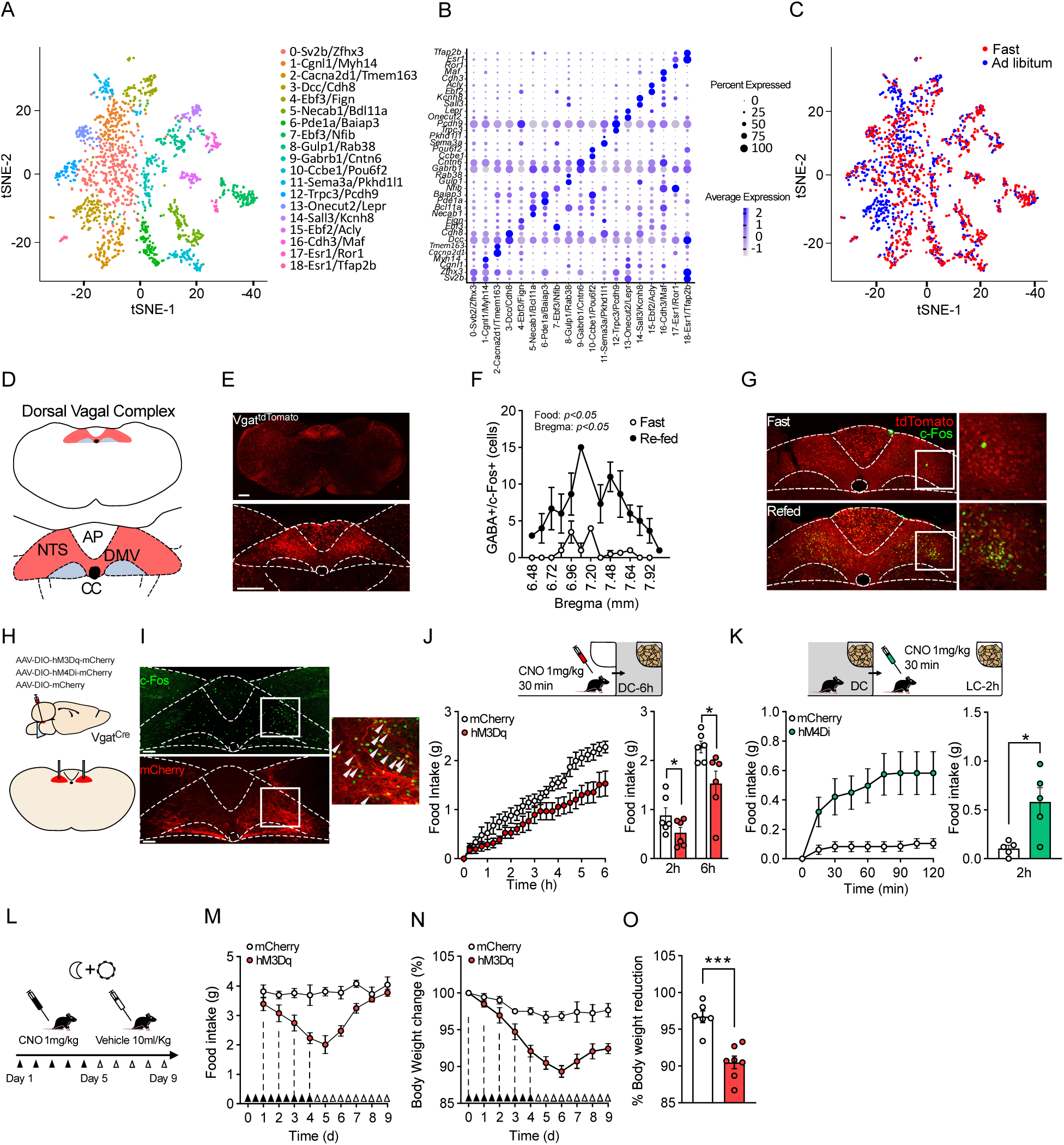
GABA^NTS^ activation reduces food intake and body weight. (A) tSNE plot of *Slc32a1^+^* neurons in the DVC colored by cluster, (B)Dot plot showing scaled expression of marker genes for each of the 19 *Slc32a1^+^* clusters (C) tSNE plot of the *Slc32a1^+^* nuclei colored by nutritional status (D) Schematic of the DVC (top) with zoom-in diagram showing AP, NTS and DMC (bottom). (E) Representative photomicrograph of GABAergic cell distribution in the medial brainstem (top) and in the DVC (bottom) using *Vgat^tdTom^* mice. (F) Quantification GABAergic cells expressing c-Fos following 16h fasting and fasting+2h refeeding (n=3, Bregma level: two-way ANOVA F_(13,34)_=2.63; p=0.012; Nutritional state: two-way ANOVA F_(1,4)_=8.69; p=0.042. (E) Representative micrograph of *Vgat^tdTom^* cells (red) expressing c-Fos (green) in mice fasted 16h (top) and re-fed for 2h (bottom). (H) Schematic of AAV-DIO-hM3Dq/hM4Di/mCherry infused into NTS of *Vgat^Cre^* mice. (I) Representative photomicrograph of c-Fos (green) expression in hM3Dq-mCherry (red)-expressing cells in the NTS of VgatCre mice treated with CNO (1 mg/kg, i.p) and (J) 6h cumulative and (J) 2h and 6h total food intake (n=6, Unpaired t test t_(11)_=2.663, p=0.0238) of GABA^NTS^:hM3Dq CNO injected mice compared to mCherry. (K) 2h cumulative and (J) 2h total food intake (n=5, Unpaired t test t_(8)_=3.205, p=0.0125) GABA^NTS^:hM4Di CNO injected mice compared to mCherry. (L) 10-day treatment protocol schematic. Daily CNO treatment significantly reduced (M) food intake (RM two-way ANOVA F_(1,11)_=20.57; P=0.0008), (N) body weight (n=6/7, RM two-way ANOVA (F_(1,11)_=21.96; p=0.0007) and (O) body weight change GABA^NTS:hM3Dq^ mice compared to control GABA^NTS^:mCherry mice (n=6/7, Unpaired t test t_(11)_=5.252, p=0.0003). C-D n=4/group, H-K n=6; M-O n=6 mCherry and n=7 hM3Dq. Data represented as mean±S.E.M. *: p<0.05; **: p<0.01; ***: p<0.001. AP: Area Postrema; CC: Centre canal; DMV: Doral motor nucleus of the vagus nerve; NTS: Nucleus of the solitary tract.

We therefore, examined whether GABA^NTS^ cells are responsive to energy status. We facilitated the visualization of GABA^NTS^ neurons by crossing *Vgat-ires-Cre* mice with *tdTomato^fl/fl^* reporter mice (*Vgat^tdTom^,* **Figure 1D and 1E**). We then examined the expression levels of the neuronal activity marker c-Fos (Olson et al., 1993) in mice that were overnight fasted and in mice that were refed for 2 hours after overnight fasting. Re-fed mice showed significantly more c-Fos in the NTS (**Figure S1D**) and specifically an increased number of GABAergic cells activated (**Figure 1F and 1G**) when compared to fasted mice. This suggests that GABA^NTS^ cells are responsive to positive energy status.

To determine whether GABA^NTS^ neurons have a role in the control of food intake, we used chemogenetic Designer Receptors Exclusively Activated by Designer Drugs (DREADD) to manipulate GABA^NTS^ cellular activity (Alexander et al., 2009). Specifically, AAVs expressing hM3Dq, hM4Di or control mCherry were bilaterally injected into the NTS of *Vgat^Cre^* mice (**Figure 1H**). This allowed to modulate GABA^NTS^ neuron activity with the administration of the designer drug clozapine-N-oxide (CNO). Administration of CNO in hM3Dq-injected mice induced strong c-Fos expression in the NTS (**Figure 1I**). We next assessed the effect of the modulation of GABA^NTS^ on food intake in mice under different scenarios. In ad libitum fed mice, hM3Dq chemogenetic activation of GABA^NTS^ cells significantly reduced overall food intake during the active dark cycle when compared to control (mCherry) siblings (**Figure 1J and S1E**). Similarly, hM3Dq GABA^NTS^ cell activation reduced food intake in hunger-induced fasted mice when compared to control mice (**Figure S1F and S1G**). Conversely, hM4Di chemogenetic inhibition of GABA^NTS^ cell in satiated mice during the light cycle significantly increased food intake when compared to satiated littermate mCherry controls (**Figure 1K**).

Given the potent reduction of food intake produced by the activation of GABA^NTS^ neurons, we examined whether prolonged activation impacted body weight. Twice-daily administration of CNO (**Figure 1L)** in hM3Dq-expressing mice induced a significant reduction in daily food intake (**Figure 1M**) and a progressive reduction in body weight (**Figure 1N and 1O**). Although feeding returned to baseline levels 72 hours after the final CNO administration, body weight remained significantly lower in hM3Dq-expressing mice (**Figure 1N**). In contrast, prolonged GABA^NTS^ neuron inhibition using hM4Di did not induce greater food intake (**Figure S1H**) nor changes in body weight (**Figure S1I**) when compared to their littermate mCherry controls. These results indicate that the selective activation of GABA^NTS^ neurons is sufficient to result in a sustained reduction in energy intake and hence body weight.

Recently, it has been reported that the inhibition of neurons expressing the GABA-producing enzyme, glutamate decarboxylase (GAD), in the NTS of rats partially blunted the anorectic effect of the GLP1-R agonist liraglutide (Fortin et al., 2020). These data, suggest that GABA^NTS^ contributes to the therapeutic effects of GLP1-R agonists in rats. However, GLP-1R expression in rats and mice differs, especially its receptor density (Cork et al., 2015; Graham et al., 2020; Jensen et al., 2018). We therefore assessed the feeding response of liraglutide in GABA^NTS^-hM4Di/mCherry-expressing mice. Chemogenetic inhibition of GABA^NTS^ partially blunted the acute anorectic effects of liraglutide (**Figure S1J and S1K**). This provides support that GABA^NTS^ participates in the anorectic effects of liraglutide in mice.

### Optogenetic stimulation of GABA^NTS^ terminals inhibits AgRP/NPY^ARC^ cells

To clarify the circuitry through which GABA^NTS^ neurons influence feeding and body weight, we used Channelrhodopsin 2 (ChR2)-assisted circuit mapping (CRACM) (González et al., 2016; Petreanu et al., 2007). Specifically, an AAV expressing ChR2-mCherry was injected into the NTS of *Vgat^Cre^* mice (**Figure 2A**) and projection patterns were analyzed. GABA^NTS^ cells project to hypothalamic subregions including the ARC, paraventricular nucleus (PVH) and dorsomedial nucleus (DMH) and various other extra-hypothalamic regions (**Figure S2A**). Within the ARC, we observed a dense array of projections (**Figure 2B**). This identified the ARC as a candidate second order region involved in the GABA^NTS^ control of food intake and body weight.

**Figure 2.**
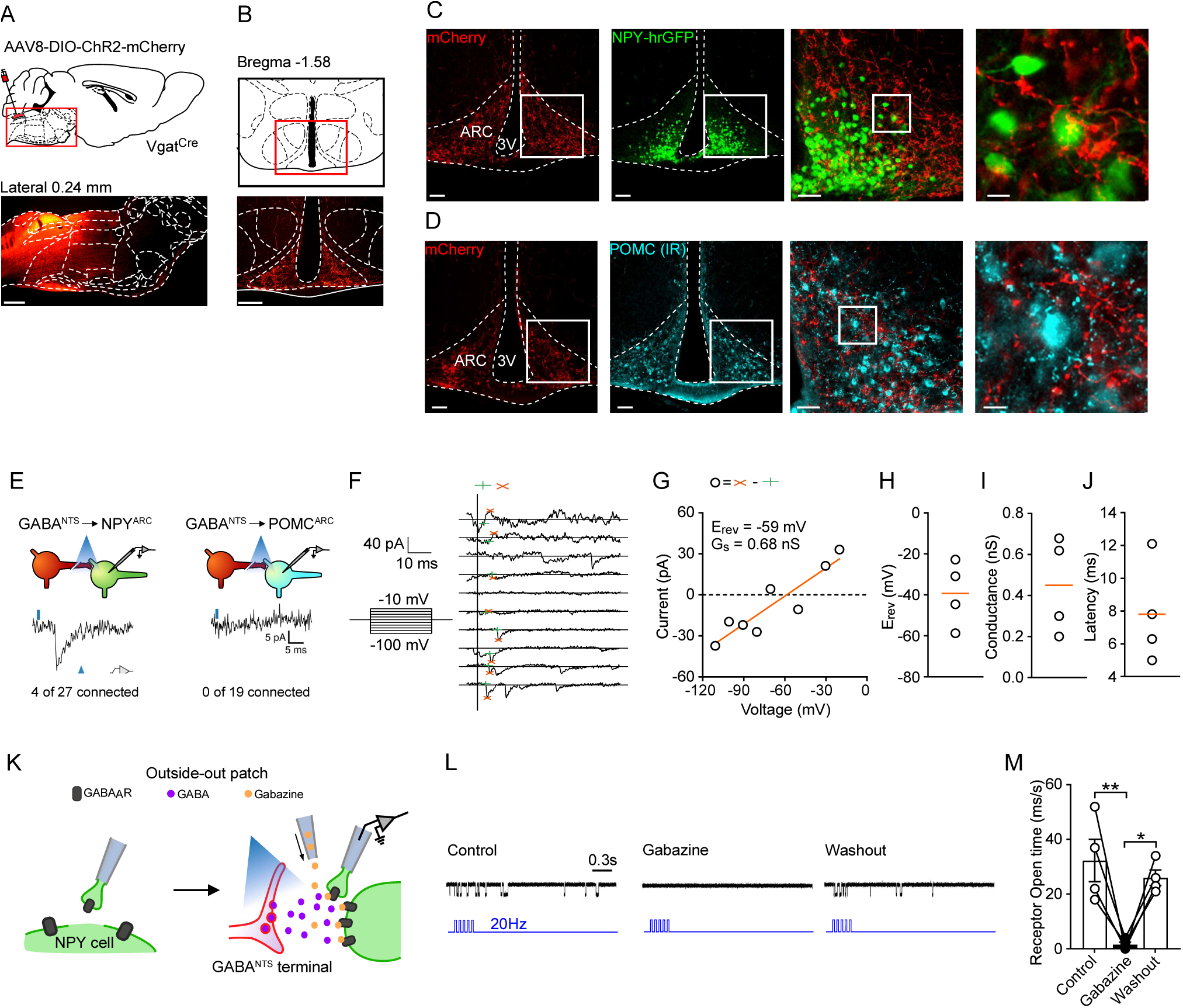
GABA^NTS^ projections to the ARC release GABA into AgRP/NPY neurons. (A) Illustration depicting details for NTS injection of AAV-DIO-ChR2-mCherry (top) and representative photomicrograph (scale 1mm) of injection (bottom) in *Vgat^Cre^* mice and (B) ChR2-mcherry containing fibers in the ARC (scale 200um). (B) Representative photomicrographs (scale 100um) and magnification (scale 50 and 10um) ChR2-mCherry fibers in the ARC close to NPY^hrGFP^-expressing cells and (D) POMC-expressing cells and in *Vgat^Cre^* mice bilaterally injected AAV-DIO-ChR2-mCherry. (E-J) *Vgat^Cre^*:*Npy^hrGFP^* mice CRACM study. (E) Representative membrane potential response of NPYGFP and POMCDsRed cells after light stimulation of mCherry containing terminals. (F) Representative membrane current in Npy^hrGFP^ cell recorded during a voltage-clamp experiment from -100mV to -10mV in 10-mV steps. (Vertical line: light pulse; green cross: base line; red cross: peak value). (G) Representative light-induced post-synaptic current during voltage clamp experiment showing peak conductance, G_s_, and reversal potential, E_rev_. (H) Individual values and median of (H) E_rev_, (I) Conductance, G_s_ and (I) latency of all responsive cells. (K) Diagram of outside-out patch technique in *Vgat^Cre^*:*Npy^hrGFP^* mice bilaterally injected AAV-DIO-ChR2-mCherry. (L) Representative channel opening recordings of control (left) gabazine (middle) and washout (right) recordings. (M) Quantification of receptor opening time (n=4, RM two-way ANOVA F_(2,6)_=16.14; p=0.0039, Bonferroni adjusted p=0.0051 Control vs Gabazine and p=0.0156 Gabazine vs Washout. Data in M are expressed as mean±S.E.M. E-J: 4/27 cells NPYhrGFP and 0/19 cells POMCdsRed; K-M: n=4-5 patches. *: p<0.05; **: p<0.01.

Fasting induces activation of AgRP/NPY-expressing neurons (Hahn et al., 1998) and direct AgRP/NPY neuron activation induces robust feeding (Aponte et al., 2011; Krashes et al., 2011). Since GABA-releasing neurons are the main inhibitory network in the brain (Krnjević & Schwartz, 1967), we hypothesized that GABA^NTS^ cells projecting to the ARC would target AgRP/NPY neurons to decrease food intake. To interrogate this potential GABA^NTS^→AgRP/NPY^ARC^ circuit, *Vgat^Cre^* were crossed with *Npy^hrGFP^* mice (*Vgat^Cre^::Npy^hrGFP^*) and bilaterally injected with AAV-ChR2-mCherry into the NTS. We observed that a subset of NPY^hrGFP^ cell bodies were surrounded by mCherry-containing fibers (**Figure 2C**). In contrast, terminals were not found surrounding neurons expressing other neuropeptides involved in the regulation of food intake such as POMC neurons (Williams & Schwartz, 2005) (**Figure 2D**).

We next investigated whether this anatomical connectivity produced functional interactions between GABA^NTS^ terminals and AgRP/NPY^ARC^ and POMC neurons. Specifically, photo-stimulation of ChR2-containing axon terminals from GABA^NTS^ neurons produced robust synaptic responses in 14% of AgRP/NPY cells in the ARC but not in POMC cells (**Figure 2E**). The rapid synaptic currents triggered in AgRP/NPY cells by the optical stimulation changed polarity near the equilibrium potential for chloride (**Figure 2F and 2G**), as expected from ionotropic GABA receptors. Light-induced currents were unexpectedly small, and their reversal potential was more positive than that expected for GABA-activated currents (about -60 mV) (**Figure 2H-J**). These effects could be explained by voltage- and space-clamp errors (Spruston et al., 1993) and in turn suggest that perhaps GABA^NTS^ terminals reach ARC NPY cells at dendrites distant from the soma. While it is not possible to directly demonstrate that this is the case, we tested whether this reasoning was justified by simulating the optogenetic activation of GABA synaptic events at distal vs proximal dendrites in a model neuron. Using a predictive neuronal model (Lindroos & Hellgren Kotaleski, 2021), we found that GABAergic post-synaptic currents become progressively smaller with their reversal potential progressively more positive, the further the GABA inputs are from the soma (**Figure S2 B-E**).

Next, we tested whether activation of GABA receptors at NPY cells is accompanied by release of GABA from GABA^NTS^ ChR2-containing terminals. We first stimulated ChR2-mCherry expressing GABA^NTS^ fibers in the ARC and performed a “sniffer patch” experiment registering GABA_A_R single-channel openings above the NPY^hrGFP^ cells contacted by GABA^NTS^ fibers (**Figure 2K**). In this experiment, the GABA_A_R response was isolated with a specific cocktail of antagonists (see Methods). This provides semi-quantitative monitoring of extracellular levels of GABA (Sylantyev et al., 2020). A burst of light directed to GABA^NTS^ ChR2-containing terminals in the ARC induced single-channel openings in membrane patch in control conditions, and the currents had amplitudes comparable to GABAergic inhibitory currents (Sylantyev et al., 2020) (**Figure 2L, *left***). Application of GABA_A_R competitive antagonist gabazine reversibly blocked the single-channel openings (**Figure 2L, *middle and right*)**. The same pattern of receptor opening time was observed in all assessed patches (**Figure 2M**). This suggested that postsynaptic NPY cells are sensitive to GABA^NTS^ presynaptic release of GABA.

To test whether NPY cells were inhibited by the release of GABA from GABA^NTS^ terminals, we designed a protocol of additive subthreshold electrical stimuli. This method is designed to evoke an action potential after 5 stimuli. In addition, we coupled a 470 nm light burst to the same trigger (**Figure S2F**). We then alternated a sequence of electrical stimuli and electrical stimuli with light burst. The electrical stimulation evoked an action potential(s) (**Figure S2G, *left*)** followed by low-frequency or no single-channel openings in the sniffer patch. When we coupled the electrical stimulation with the light burst, the occurrence of actions potentials was blocked and accompanied with single-channel openings in a sniffer patch (**Figure S2G, *right***). This happened in all cells patched (**Figure S2H**). These openings resembled GABA_A_R openings illustrated in Figure 2L. Subsequent stimulations showed a decrease in channel opening intensity, suggesting a depletion of GABA stores from the presynaptic terminal, which was eventually insufficient to prevent the action potential (**Figure S2I**). These results provide strong evidence that GABA^NTS^ fibers inhibit AgRP/NPY cells due to the release of the fast neurotransmitter GABA.

### GABA^NTS^→ARC optogenetic stimulation reduces feeding and is not aversive

Given the dense fiber projection pattern of NTS-GABAergic cells to the ARC, and that its activation induced a strong inhibition of AgRP/NPY neurons, we interrogated whether this circuit is sufficient to influence food intake. To investigate this, AAVs expressing ChR2-mCherry were infused into the NTS of *Vgat^Cre^* mice and an optic fiber was placed above the ARC (**Figure 3A**). Food intake was measured both without and with ARC photo-stimulation prior to the onset of the dark cycle (**Figure 3B and 3C**). Light stimulation of GABA^NTS^→ARC terminals induced a strong acute inhibition of food intake that lasted 60 min from food presentation, as compared to the same mice without light stimulation (**Figure 3D and 3E**).

**Figure 3.**
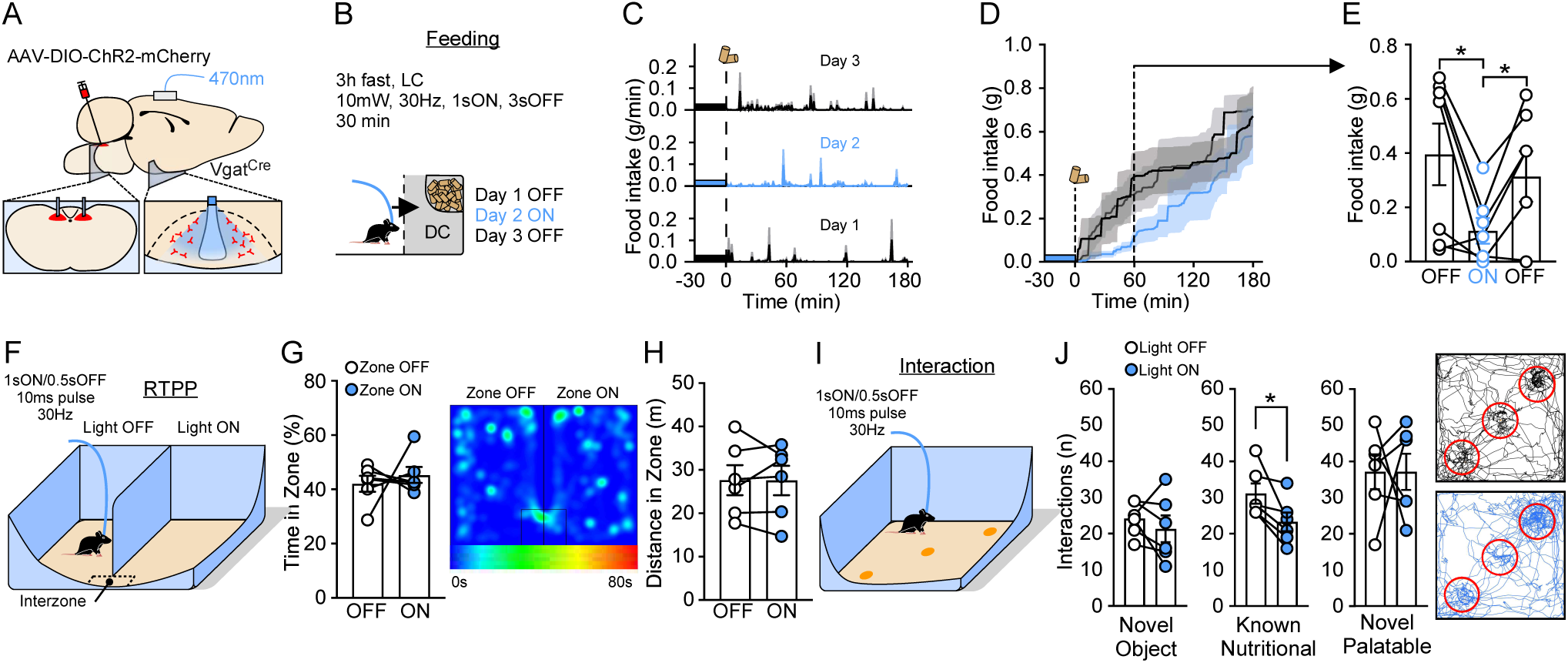
Activation of GABA^NTS^→ARC projections reduces food intake without inducing aversion. (A) Schematic of viral infection and optic fiber placement for in vivo stimulation of GABA^NTS^→ARC. (B) Protocol of stimulation. (C) Representative 1min resolution time plot of 3h food intake measurement. GABA^NTS^→ARC stimulation significantly reduces (D) cumulative 3h food intake and (E) 60min intake (n=7, day 1 (OFF) vs day 2 (ON) paired Student t-test t_(6)_=3.035, p=0.0229; day 2 (ON) vs day 3 (OFF) paired Student t-test t_(6)_=3.283, p=0.0168). (F) Diagram illustrating RTPP task. GABA^NTS^→ARC does not alter (G) time spent (representative heat map of time) or (H) distance travelled in stimulated and non-stimulated zones. (I) Diagram illustrating an object interaction task. (J) Interactions with novel, known nutritional or novel palatable objects and a representative path track around each object (red circle). GABA^NTS^→ARC stimulation reduced interaction with known nutritional item (n=6, Student’s t test t_(5)_=3.043, p=0.0287). Data are expressed as individual values and as mean±S.E.M, n=6 mice. *: p<0.05; RTPP: real-time place preference.

The DVC has been proposed to be a key region modulating food intake reduction associated with aversive states (D’Agostino & Luckman, 2022). To assess whether GABA^NTS^→ARC activation produces aversion or negative valence (Berridge, 2004; Betley et al., 2015), we evaluated the existence of passive avoidance behavior using an adapted real-time place preference (RTPP) task (D’Agostino et al., 2016; Kim et al., 2013; Stamatakis & Stuber, 2012). Specifically, in a two-sided open arena, light stimulus was coupled to one of the sides (**Figure 3F**). GABA^NTS^→ARC stimulation or lack of stimulation did not produce a place preference (**Figure 3G**) and mice travelled a similar distance in both sides of the arena (**Figure 3H**). These findings indicate that GABA^NTS^→ARC stimulation does not produce negative valence or aversion.

In addition to influencing homeostatic feeding, modulation of AgRP/NPY cells has also been reported to elicit anxiety-like behaviors influencing exploration and foraging which impacts food consumption (Dietrich et al., 2015; Heinz et al., 2021; Li et al., 2019). To test whether the activation of the GABA^NTS^→ARC circuit produced anxiety-related behavior, mice were assessed in an open-field arena (OFA) and an elevated zero-maze task (EZM) (**Figure S2A and S2D**). GABA^NTS^→ARC stimulation did not alter the time mice spent in the center of the OFA (**Figure S2B**). Likewise, mice displayed similar ambulatory patterns and travelled comparable distance during the OFA test with and without GABA^NTS^→ARC stimulation (**Figure S2C**). Consistent with the OFA data, GABA^NTS^→ARC stimulation did not alter either the time (**Figure S2E**) or distance travelled in the exposed zones of the EZM (**Figure S2F**). These results provide evidence that stimulation of the GABA^NTS^→ARC circuit impacts feeding without altering other behavioral states such as anxiety-like behavior.

We postulated that since our model involved fast neurotransmission rather than long term release of neuropeptides, the reduction in food intake that we observed could begin by reducing the interaction with the nutritional cue at first instance. To investigate this, we designed a task where mice would first be allowed to explore an empty arena and then three items would be presented at the same time, an inedible object, a novel palatable food item and a known nutritional food item (**Figure 3I**). The number of interactions with each item was measured with and without GABA^NTS^→ARC stimulation. GABA^NTS^→ARC stimulation did not induce changes in the interaction with the novel object (**Figure 3J**), further suggesting that GABA^NTS^→ARC does not induce anxiety-like behavior. The number of interactions with the novel palatable food item was increased under both non-stimulation and stimulation trials (**Figure S2F and S2G**). However, we found that under optical stimulation mice had less interactions with the known nutritional item (**Figure 3J and S2G**), suggesting that GABA^NTS^→ARC activation reduces hunger. Taken together, these findings indicate that GABA^NTS^→ARC stimulation decreases feeding and hunger without inducing aversion or anxiety.

### GABA^NTS^→ARC neuron activation promotes satiety

We next used a two-virus intersectional approach (Fenno et al., 2014, 2017) to provide a more detailed characterization of GABA^NTS^→ARC activation in energy homeostasis. *Vgat^Cre^* mice were bilaterally injected with AAVs expressing Flipase recombinase (FlpO) (AAV-DIO-FlpO) under the control of Cre into the NTS of *Vgat^Cre^* mice, allowing us to express a second recombinase only in GABA^NTS^ cells. After surgery recovery, a retrograde AAV encoding for a FlpO dependent hM3Dq-mCherry (rgAAV-fDIO-hM3Dq-mCherry) was bilaterally injected into the ARC to retrogradely deliver hM3Dq in a FlpO-dependent manner to GABA^NTS^ cells. Therefore, hM3Dq was expressed only in GABA^NTS^ cells projecting to the ARC in *Vgat^Cre^* mice (**Figure 4A**).

**Figure 4.**
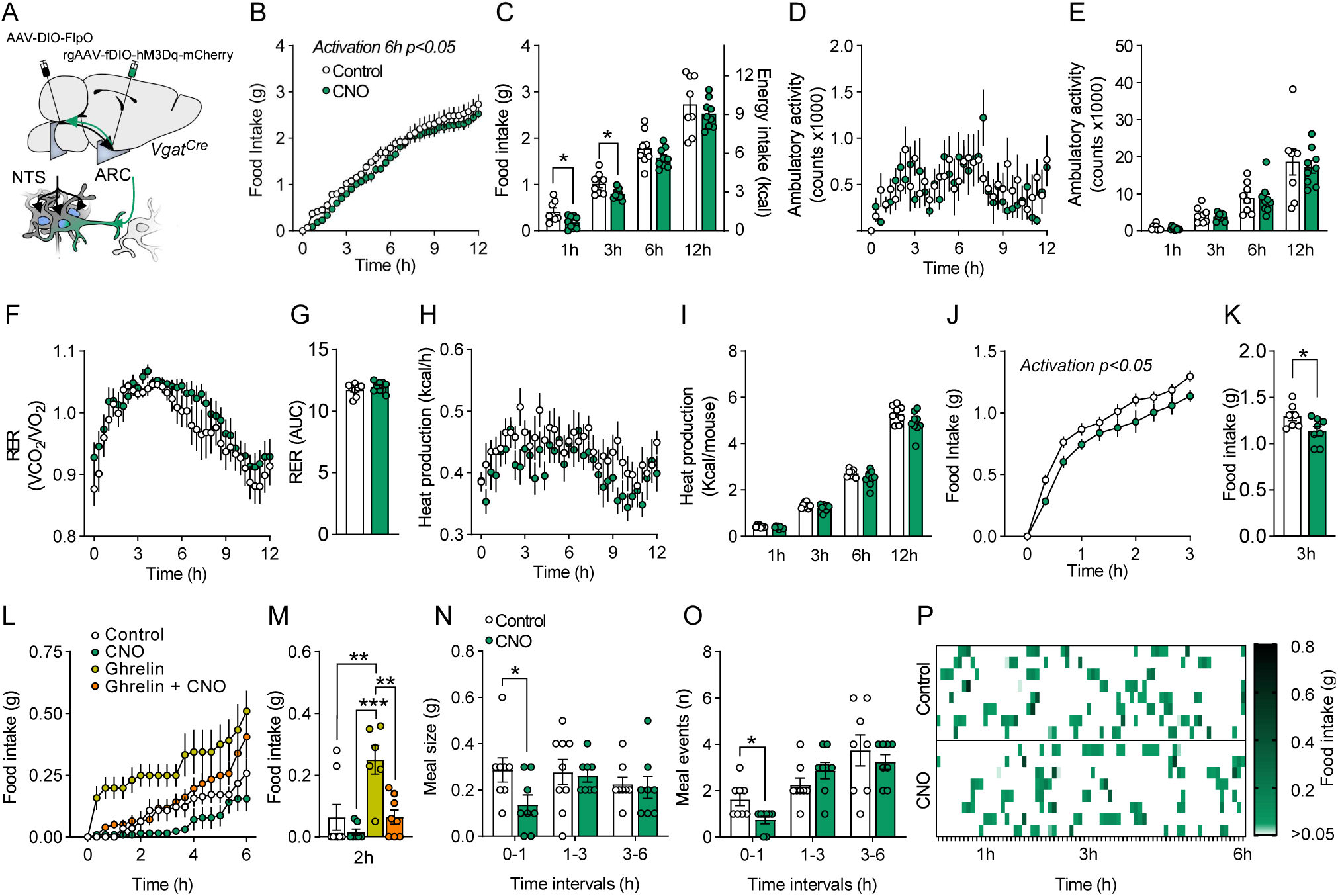
Chemogenetic activation of GABA^NTS^→ARC neurons induces satiety. (A) Diagram illustrating the two-virus intersectional strategy to express hM3Dq only in GABA^NTS^ neurons projecting to the ARC. Mice were treated with saline or CNO (1 mg/kg, i.p.). (B) CNO significantly reduced cumulative food intake over 6h (0-6h F_(1,15)_=7.418, p=0.0157), (C) 1h (Unpaired t-test t_(15)_=2.648, p=0.0183) and 3h (Unpaired t-test, t_(15)_=2.568 P=0.0214) food intake compared to saline. (D) CNO did not alter 12h or (E) 1, 3, 6 or 12h quantification of ambulation; (F) 12h RER or (G) AUC quantification; (H) 12h or (I) 1, 3, 6 and 12h heat production per mouse compared to saline. (J-K) CNO significantly reduced 3h food intake following overnight fasting (J, 2-way ANOVA F_(1.13)_=6.139, p=0.0277 and K, unpaired t-test t_(13)_=2.320, p=0.0372). (L-M) CNO attenuated ghrelin hyperphagia over 2h (RM ANOVA F_(3,16)_=11.12; p=0.0003, Bonferroni adjusted p=0.0042 control vs ghrelin, p=0.0003 CNO vs ghrelin and p=0.002 Ghrelin vs Ghrelin+CNO). (N) CNO reduced meal size (Unpaired t-test, t_(14)_=2.256, p=0.0406) and (O-P) decreased meal number (Unpaired t-test, t(14)=2.824, p=0.0135) compared to control. Data are expressed as individual values and as mean±SEM. *: p<0.05; **: p<0.01. ***: p<0.001.

Corroborating the optogenetic data presented above, the selective chemogenetic stimulation of the GABA^NTS^→ARC circuit significantly reduced acute food intake (**Figure 4B** and **4C**) in ad libitum fed mice. GABA^NTS^→ARC circuit activation did not alter overall locomotor activity (**Figure 4D** and **4E**), respiratory quotient (**Figure 4F** and **4G**) or heat production (**Figure 4H** and **4I**). GABA^NTS^→ARC neuron activation also reduced food intake in refed mice after overnight fasting (**Figure 4J**). Fasting is associated with a rise in systemic levels of ghrelin (Cummings et al., 2001) which acts as a pre-prandial effector stimulating AgRP/NPY neurons to initiate a feeding response (H. Y. Chen et al., 2004; Luquet et al., 2007). To examine whether GABA^NTS^→ARC neuron activation is sufficient to dampen a hunger cue, GABA^NTS^:hM3Dq-expressing mice were pre-treated with CNO prior to ghrelin. Activation of the GABA^NTS^→ARC neurons with CNO prevented the feeding induced by an orexigenic dose of ghrelin (**Figure 4L and 4M)**. We next performed an analysis of the microstructure of the feeding event during the anorectic episode produced by CNO (Clifton, 2000; Richard et al., 2011; Zorrilla et al., 2005). GABA^NTS^→ARC neuron activation reduced meal size (**Figure 4N**) and significantly reduced the number of meal events in the earliest interval compared to saline treatment (**Figure 4O** and **4P**). Taken together, these findings indicate that GABA^NTS^→ARC neuron activation is sufficient to blunt fasting and ghrelin-induced hunger and significantly reduces food intake by promoting satiety.

## Discussion

Here we identify a critical new brain circuit modulating appetite and body weight. We focused on the NTS because it is a brain region positioned to receive and integrate energy-related information from the periphery and relay it within the CNS to promote energy homeostasis. However, the NTS is neurochemically heterogeneous and key neurons within it performing this function have not been fully defined. Using multi-methodological approach, here we identify GABA^NTS^ neurons as sufficient to control feeding behavior and body weight in mice.

Recent efforts to decode the function of specific chemically defined neurons within the NTS have revealed that distinct subpopulations of glutamatergic cells play a role in energy homeostasis (Dowsett et al., 2021). However, NTS inhibitory GABA-releasing neurons have not been studied in detail and this is necessary to clarify the role of both excitatory and inhibitory NTS signals in the regulation of energy balance (Cheng et al., 2022). Here we provide a detailed characterization of the effect of GABA^NTS^ in the regulation of energy homeostasis and body weight. A recent report provided evidence that obesity medication liraglutide engages GABA^NTS^ neurons to reduce food intake in rats, providing a rationale that activating GABA^NTS^ cells may have translational relevance for the treatment of human obesity (Fortin et al., 2020).

The NTS is involved in satiety and satiation, and refeeding induces a strong neuronal activation in this region (D’Agostino et al., 2016; Wu et al., 2014). We discovered that refeeding significantly activates a subset of GABA^NTS^ cells. Further, we found that activation of GABA^NTS^ neurons with chemogenetics was anorectic, and controlled food intake and body weight. Consistently, when we inhibited these neurons, we observed a potent induction of feeding in satiated mice, suggesting a crucial role in the induction of early eating. However, when the inhibition occurred in ad libitum mice entering the dark cycle, we didn’t observe an increased food intake. This could be explained by two reasons. First, despite of mice being kept in ad libitum conditions, they are not satiated at the onset of the dark cycle, meaning GABA^NTS^ neuros are likely to not be active at this time, therefore an inhibition would not meaningful. Second, during the active dark cycle there are multiple feeding cues not only vagal sensory afferents but afferent from the sensory system that would modulate feeding and could counteract GABA^NTS^ lack of inhibition in second order neurons. Our data illustrates that GABA^NTS^ neurons project widely within the brain, with particularly dense innervation of the hypothalamus. Several studies indicate that there is a coordination between the NTS and the hypothalamus to orchestrate the meal event (Aklan et al., 2020; Blevins et al., 2004; D’Agostino et al., 2018; Liu et al., 2017; Tsang et al., 2020). However, whether the ARC receives inhibitory control from the NTS is not known and was examined here.

Given that GABA is an inhibitory neurotransmitter, we hypothesized that GABA^NTS^ neurons inhibit appetite stimulating neurons. We focused on AgRP/NPY^ARC^ neurons because of the dense GABA^NTS^ innervation that we found and the potent orexigenic properties of AgRP/NPY (Aponte et al., 2011; Krashes et al., 2011; Luquet et al., 2005). In addition, AgRP/NPY neurons display a multiple timescale activation (Mandelblat-Cerf et al., 2015) that could explain the lack of feeding effect during the chemogenetic inhibition of GABA^NTS^. We demonstrated that GABA^NTS^ terminals in the ARC release GABA and that this is synchronized with suppression of the action potential propagation postsynaptic AgRP/NPY^ARC^ cells. The timescale of GABA_A_R openings after optical stimulation in a membrane patch placed over the GABA^NTS^ terminal suggests the release of GABA from this terminal rather than from another inhibitory neurons. The number of AgRP/NPY neurons responding is in line with the 20% ARC responders to NTS innervation published previously (Aklan et al., 2020). However, despite the relatively small number of AgRP/NPY^ARC^ neurons inhibited by GABA^NTS^ terminals, it was sufficient to significantly reduce feeding in freely behaving mice. AgRP/NPY cells are poised to orchestrate the integration of homeostatic, reward and sensory cues as well as learning and conditioned behaviors (Aponte et al., 2011; Berrios et al., 2021; Deem et al., 2022; Dietrich et al., 2012, 2015; Garau et al., 2020; Han et al., 2021; Jikomes et al., 2016; Krashes et al., 2011, 2014; Wang et al., 2021). A detailed analysis of feeding behavior revealed that activation of the GABA^NTS^→ARC pathway specifically reduces hunger and promotes satiety and does not induce negative valence, aversion or anxiety.

Obesity is an international health concern that is primarily the consequence of over-eating. Defining the mechanisms governing hunger and food intake is therefore of paramount importance. Here we identify a new player that controls appetite and body weight, GABA^NTS^. Specifically, we show that NTS GABA-releasing neurons are active during satiety and reduce food intake and body weight without causing aversion or anxiety. We demonstrate that GABA^NTS^ cells directly activate GABA_A_Rs on the surface of AgRP/NPY cells which inhibits neuron activity. These studies reveal for the first time the effect of GABA^NTS^ neurons on feeding and body weight, and identify a fast inhibitory circuit between the NTS and the ARC in the control of food intake. These results thereby provide significant insight into the brain circuits governing appetite and body weight, findings of relevance to the global obesity crisis.

## Methods

### RNA-Seq

Single nucleus RNA-sequencing data from the mouse hindbrain in the fed and fasted state was taken from (Dowsett et al., 2021). Neuronal nuclei expressing at least 1 UMI count for *Slc32a1* were identified as GABAergic neurons, subsetted and reclustered using Seurat package version 4.3 (Hao et al., 2021). Marker genes for each cluster were calculated using Wilcoxon’s rank-sum test. Each cluster was named with 2 marker genes that were expressed in >60% of the cluster, <30% of the rest of the data and had an average log fold change >0.5. If no genes fit these criteria, then the two genes with the lowest p-values were used. Differential gene expression analysis between ad libitum fed and overnight fasted cells was performed using the Wilcoxon’s rank sum test. Feature plots were drawn using the Seurat package and ggplot2.

### Animals

*Vgat-ires-Cre* (Vong et al., 2011) (Slc32a1tm2(cre)Lowl; #016962), *NPY-hrGFP* (van den Pol et al., 2009)(van den Pol et al., 2009)(van den Pol et al., 2009)(van den Pol et al., 2009) (B6.FVB-Tg(Npy-hrGFP)1Lowl/J; #006417), POMC-dsRed (Hentges et al., 2009) (Tg(Pomc-DsRed)18Low) and *Rosa26tdTomato-LoxP* (Madisen et al., 2009) (B6.Cg-Gt(ROSA)26Sortm9(CAG-tdTomato)Hze/J, #007909) mice were obtained from The Jackson Laboratory (Bar Harbor, USA) and bred in a C57Bl/6J background. Mice were fed with standard laboratory chow (Standard CRM (P) 801722, Special diets, UK) and provided with water ad libitum, unless otherwise stated. Mice were kept in a 12-hours light:dark cycle (7am-7pm) in environmental controlled conditions (20-22°C and 40-60% RH). All experimental procedures were performed in accordance with the UK Animal (Scientific Procedures) Act 1986.

### Viral vectors

Cre-dependent viral vectors were purchased from Addgene, AAV8-hSyn-DIO-hM3D(Gq)-mCherry (1.83x10^12^ gc/ml) was a gift from Bryan Roth (Addgene plasmid # 44361) (Krashes et al., 2011), AAVrg-hSyn-fDIO-hM3D(Gq)-mCherry-WPREpA (1.8x10^12^ gc/ml) was a gift from Ulrik Gether (Addgene plasmid # 154868); AAV8-hSyn-DIO-mCherry (3.6x10^12^ gc/ml) was a gift from Bryan Roth (Addgene plasmid # 50459); AAV8-pEF1a-DIO-FLPo-WPRE-hGHpA (2x10^12^ gc/ml) was a gift from Li Zhang (Addgene plasmid # 87306) (Zingg et al., 2017). AAV2-EF1a-DIO-ChR2(E123T/T159C)-mCherry and AAV2-EF1a-DIO-ChR2(E123T/T159C)-YFP (7.3x10^12^ vp/ml) were a gift from Karl Deisseroth and were obtained from University of North Carolina Vector Core (Chapel Hil, NC, USA). All viral particles were delivered into nuclei-specific regions through stereotaxic injections.

### Stereotaxic surgeries

For viral delivering in the NTS, stereotaxic surgery was adapted from previous studies (D’Agostino et al., 2016). Briefly, 12-20 weeks old mice were anaesthetized with isoflurane, back region of the neck shaved and placed in a stereotaxic instrument (David Kopf instruments, CA, USA) with a face mask (World Precision Instruments, FL, USA). Head was inclined ∼70 degrees forward and a longitudinal incision was made in the skin at the level of the C1; neck muscles were retracted to expose the atlanto-occipital membrane. This was carefully dissected allowing access to the dorsal brainstem and visualization of the obex. Using a pulled glass capillary (40µ tip diameter) (G1, Narishige, UK) and a pneumatic microinjector (IM-11-2, Narishige, UK) 200-300 nl of viral preparation was bilaterally injected into the NTS (obex: AP:0.25 mm AP, L:± 0.25 mm and DV:-0.25mm) at a flow of 50nl/min. Capillary was left in the injection place for 5 min to allow diffusion and it was removed slowly to avoid dispersion to neighbor brainstem regions. Viral delivery into the ARC was performed as previously described (Wagner et al., 2022) at coordinates bregma: AP:1.58 mm AP, L:±0.2mm and DV:5.90 mm. For optical fiber cannula placement, mice were allowed 4 weeks recovery form the NTS surgery and a 200 µm core diameter, 0.39NA (CFMLC, Thorlabs, UK) optical fiber implants were placed in the third ventricle above the ARC. Mice were allowed 3 weeks before any study to allow surgery recovery and maximal viral expression. Post hoc analysis of injection site, viral expression and canula placement were used as exclusion criteria for data analysis.

### In vivo photo-stimulation protocol

Optical fiber implants were attached to optogenetics patch cables (M83L1, Thorlabs, UK) connected to a rotary joint (Doric lenses) coupled to a 473-nm laser (Laserglow, Toronto, Canada) controlled via TTL-USB interface with Arduino board. For feeding experiments, the stimulation protocol was 1 s followed by 4 s break with 10ms light pulses with a frequency of 30Hz. For behavioral experiments, the stimulation protocol was 1 s followed by 0.5 s break with 10ms light pulses with a frequency of 30Hz. We used 15mW of laser power to achieve an irradiance of 5-10mW/mm^2^ (PM100D, Thorlabs) on the target area following https://web.stanford.edu/group/dlab/cgi-bin/graph/chart.php, above ChR2 threshold activation (Lin et al., 2009).

### Food intake and body weight studies

For food intake, body weight and metabolic parameters measurements, mice were single housed and habituated in indirect calorimetry system cages for one week (Phenomaster, TSE Systems, Germany). For acute ad libitum studies, access to food was removed in fed mice 2h before entering the dark cycle and CNO 1 mg/kg was i.p. administered 30 min before the dark cycle onset when food was provided. For re-feeding studies, 12 hours dark cycle-food deprived mice were i.p. injected with CNO at the beginning of the light cycle and 30 min after food was provided. For subchronic studies, mice were i.p. injected twice a day (am and pm) for 5 days with CNO following 5 days with saline.

### Behavioral tests

For valence studies, mice were assessed in an adapted real-time place preference task consisting in an open field arena with two connected identical chambers (30x25cm) (D’Agostino et al., 2016; Kim et al., 2013; Stamatakis & Stuber, 2012), one of them paired with optogenetic stimulation where mice were allowed free movement for 20 min. For anxiety tests, mice were placed in an open arena (50x50cm) with virtual delimited central and peripheral regions and allowed free movement for 10 minutes with and without stimulation in different days. For anxiety and fear assessment, mice were placed in an elevated zero maze (diameter 50cm, elevation 70 cm) with 2 hidden and 2 exposed zones and allowed free movement between zones for 10 min. Tests were performed for each animal with and without stimulation in different days. Time and locomotor parameters for each task and zone were recorded using Any-Maze software (Stoelting, IL, USA).

### Immunohistochemistry and imaging

All mice were injected with a terminal dose of anesthesia and transcardially perfused with phosphate-buffered saline (PBS) followed by 10% neutral buffered formalin. Brains were dissected, post-fixed 12 hours in formalin at 4°C, cryoprotected 48 hours with 30% sucrose 4°C and coronally sectioned in 5 series at 25 µm using a freezing microtome (8000, Bright Instruments, UK). Sections were kept in protective anti-freeze solution at 4°C until they were processed for immunohistochemistry as previously described (Yavari et al., 2016). Briefly, NTS sections were washed with PBS-0.2% Tween20 30 min and then PBS (3x10 min), blocked with 1%BSA/5%DS/0.25%Triton X-100 1 hour at room temperature and incubated with primary antibody in blocking solution with anti-c-Fos (1:2500, 2250, CST, USA), anti-c- Fos (for chromogenic) (1:5000, ABE457,Merck, UK), anti-mCherry (1:2000, AB0040-200, Scigen, PT), anti-RFP (1:1000, 600-401-379, Rockland Immunochemicals, USA), anti-POMC (1:3000, H-029-30, Phoenix Pharmaceuticals, USA), anti-hrGFP (1:2000, 240141, Agilent, USA), anti-TH (1:2000, MAB318, Merck, Germany) 16 hours at room temperature. The next day, sections were washed with PBS-Tween and PBS and incubated 1 hour with appropriate secondary antibodies in blocking solution (1:500, AlexaFluor594, AlexaFluor488, Invitrogen, UK) at room temperature. For c-Fos expression quantification in fast vs refed study, chromogenic staining with DAB reagent was performed as previously described (D’Agostino et al., 2016).

Images were acquired using Axioskope2 microscope and Axiovision software (Zeiss, Germany). All images were converted to 8-bit, peudorecolored and cells counted using ImageJ (Fiji).

### Electrophysiology

#### CRACM study

CRACM experiments were performed as previously described (González et al., 2016). Six *Vgat^Cre^*:*Npy^hrGFP^* mice and six *Vgat^Cre^*:*Pomc^dsRed^* mice bilaterally injected with AAV-ChR2-mCherry and AAV-ChR2-mCherry respectively into the NTS, aged between 5 and 7 months at the time of the electrophysiology experiments, were used. Expression of mCherry-ChR2 was targeted to NTS^GABA^ cells by stereotaxic injection of AAV as above. Coronal brain sections 180-µm thick were prepared from these mice at least 8 days after virus injections and were placed in a bath solution consisting of (in mM) 125 NaCl, 2.5 KCl, 1.2 NaH_2_PO_4_, 21 NaHCO_3_, 1 glucose, 2 MgCl_2_, 2 CaCl_2_. GFP-expressing cells in the ARC were identified using an upright microscope (Scientifica S-Scope-II) equipped with the appropriate fluorescence filters. Whole-cell recordings from these cells were obtained with glass pipettes (World Precision Instruments 1B150F-4) filled with a solution containing (in mM) 120 K-gluconate, 10 HEPES, 10 KCl, 1 EGTA, 2 MgCl2, 4K2ATP, and 1 Na2ATP, tip resistance 3-7 MOhm. Data was acquired using Axon Instruments hardware (MultiClamp 700B, Digidata 1550). To test for GABA inputs to AgRP/NPY^ARC^ cells, the membrane potential in these cells was clamped at increasing levels of voltage (from -100 to -10 mV in 10-mV increments), while ChR2-expressing terminals were stimulated by a single light pulse (CoolLED pE-4000) to induce post-synaptic currents. Liquid junction potential, estimated to be 10 mV, was subtracted from the measurements. Chloride equilibrium potential was calculated to be -60.3 mV.

#### Sniffer-patch recordings

Transverse hypothalamic slices from *Vgat^Cre^*:*Npy^hrGFP^* mice bilaterally injected AAV-DIO-ChR2-mCherry were cut at 200-250 using a Leica VT1200S vibratome. Slices were incubated for one hour in a solution containing (in mM): 124 NaCl, 3 KCl, 1 CaCl_2_, 3 MgCl_2_, 26 NaHCO_3_, 1.25 NaH_2_PO_4_, 10 D-glucose, and bubbled with 95/5% O_2_/CO_2_, pH 7.4. After incubation, slices were transferred to a recording chamber continuously superfused with an external solution. The external solution composition differed from incubation solution in containing 2 mM CaCl_2_ and 2 mM MgCl_2_.

In all experiments the intracellular pipette solution for voltage-clamp recordings contained (mM): 117.5 Cs-gluconate, 17.5 CsCl, 10 KOH-HEPES, 10 BAPTA, 8 NaCl, 5 QX-314, 2 Mg-ATP, 0.3 GTP; for current-clamp recordings: 126 K-gluconate, 4 NaCl, 5 HEPES, 15 glucose, 1 MgSO_4_·7H_2_O, 2 BAPTA, 3 Mg-ATP (pH 7.2, 295-310 mOsm in both cases); pipette resistance was 7-9 MOhm; recordings were performed at 33-35°C using Multiclamp-700B amplifier with -60 or -70 mV holding current (for voltage-clamp recordings); signals were pre-filtered and digitized at 10 kHz. In experiments where transmembrane currents were recorded in outside-out patches only (sniffer-patch recordings), the GABA_A_ receptors response was isolated with a ligands cocktail containing 50 µM APV, 20 µM NBQX, 50 nM CGP-55845, 200 µM S-MCPG, 10 µM MDL-72222, and 1 µM strychnine.

### Statistics and data analysis

Statistical analyses were performed using GraphPad Prism 9 software and are described in the figure legend where 2-tail paired and unpaired Student t-test were used when comparing 2 groups and RM/two-way ANOVA test with Bonferroni post-hoc correction when comparing 4 groups. RNA-Seq data were analyzed as described above. No statistical method was used to predefine sample size, randomization and blinding was performed for histological quantifications. Statistical significance was accepted when the p≤ 0.05. Raw data was stored in Excel and figures assembled with CorelVector and Affinity Designer software.

## Author Contributions

The project was conceived by PBM and LKH. PBM designed and performed experiments and analyzed data with assistance from YM; GKCD and GSHY performed and analyzed the RNA-Seq data. SS and AG designed and performed electrophysiological recordings. The manuscript was drafted by PBM with input from all other authors.

## Acknowledgements

We gratefully acknowledge Dr F. Naneix for advice on optogenetics and editorial advice, and staff within the University of Aberdeen Medical Research Facility and the Microscopy Facility for their technical assistance. This work was supported by the ERC (MSCA-IF-NeuroEE-660219) to PBM, Wellcome Trust Institutional Strategic Support Fund (204815/Z/16/Z) to PBM and LKH, and the Biotechnology and Biological Sciences Research Council (BB/V010557/1) to JAG and (BB/V016849/1) to LKH and SS. GKCD is funded by a BBSRC CASE 4-year PhD studentship, co-funded by Novo Nordisk. GSHY is funded by the UK Medical Research Council (MC_UU_00014/1).

---

**Supplementary Figure 1.**
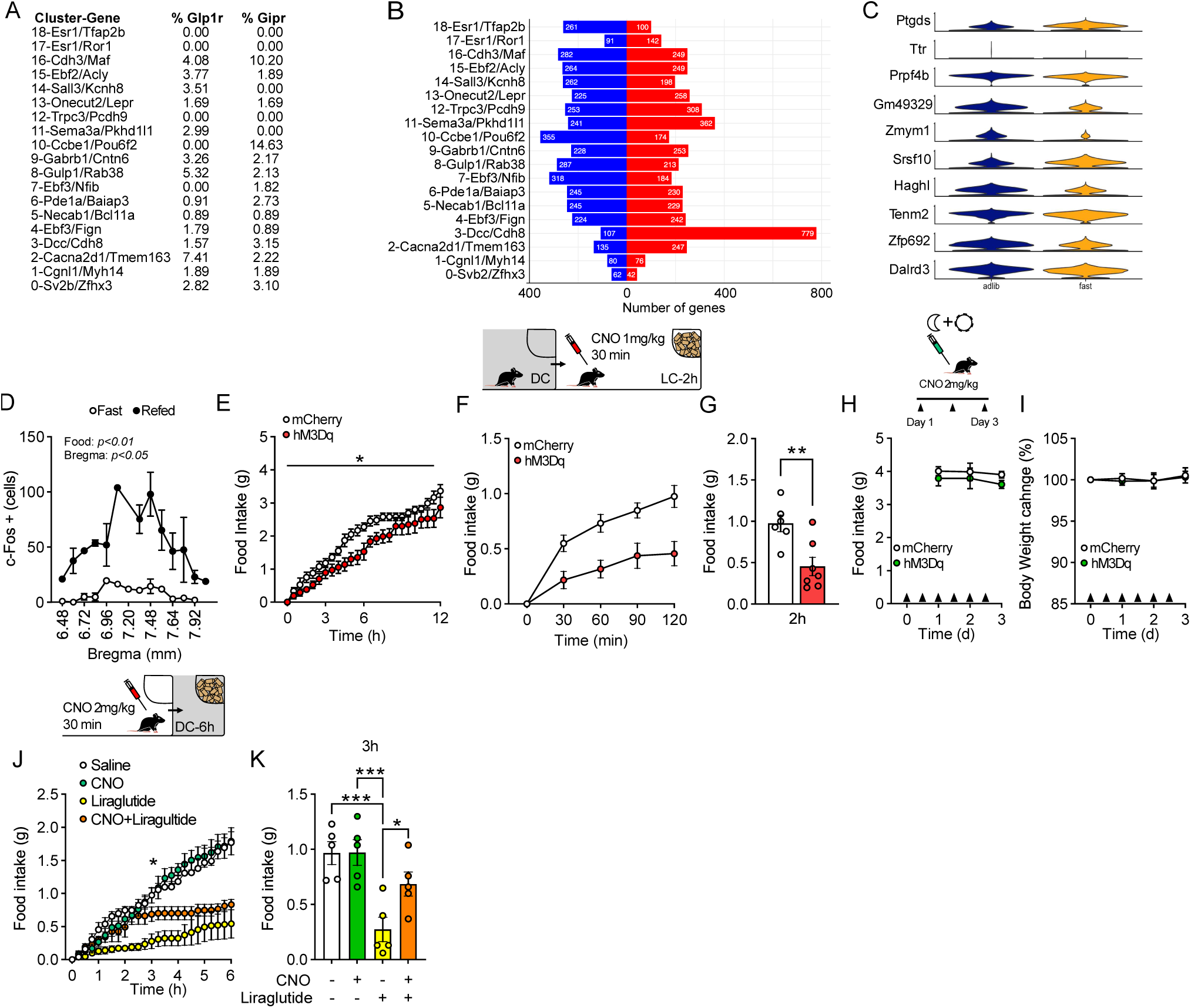
(A) Proportion of GLP1-R and GIPR expression in each *Slc32a1^+^* cluster (B) Number of genes upregulated (red) and downregulated (blue) in response to an overnight fast in each cluster. Genes included were significantly differentially regulated (P < 0.05). (C) Top 10 genes differentially expressed in ad libitum vs fasting in *Slc32a1^+^* neurons (D) Quantification of total NTS c-Fos-expressing cells following 16h fasting and fasting+2h refeeding (n=3, Bregma level: two-way ANOVA F_(13,34)_=3.16; p=0.004; Nutritional state: two-way ANOVA F_(1,4)_=18.61; P=0.012). (E) 12h dark cycle food intake in hM3Dq vs mCherry mice injected with CNO. (F) Food intake following an overnight fast compared to mCherry in GABA^NTS^:hM3Dq mice (n=6, RM two-way ANOVA F_(1,11)_=10.97; P=0.0069) and (G) 2h food intake (n=6/7, Unpaired t test, t_(11)_=3.392 p=0.006). (G-I) Food intake and body weight change in hM4Di vs mCherry mice twice daily injected with CNO (2mg/Kg). (J-K) Food intake in hM4Di mice co-administered with CNO and Liraglutide (RM: ANOVA F_(3,12)_=15.64; P=0.002, Bonferroni adjusted p=0.0004 Saline vs Liraglutide, P=0.0004 CNO vs Liraglutide and P=0.0263 Liraglutide vs Liraglutide+CNO).

**Figure S2.**
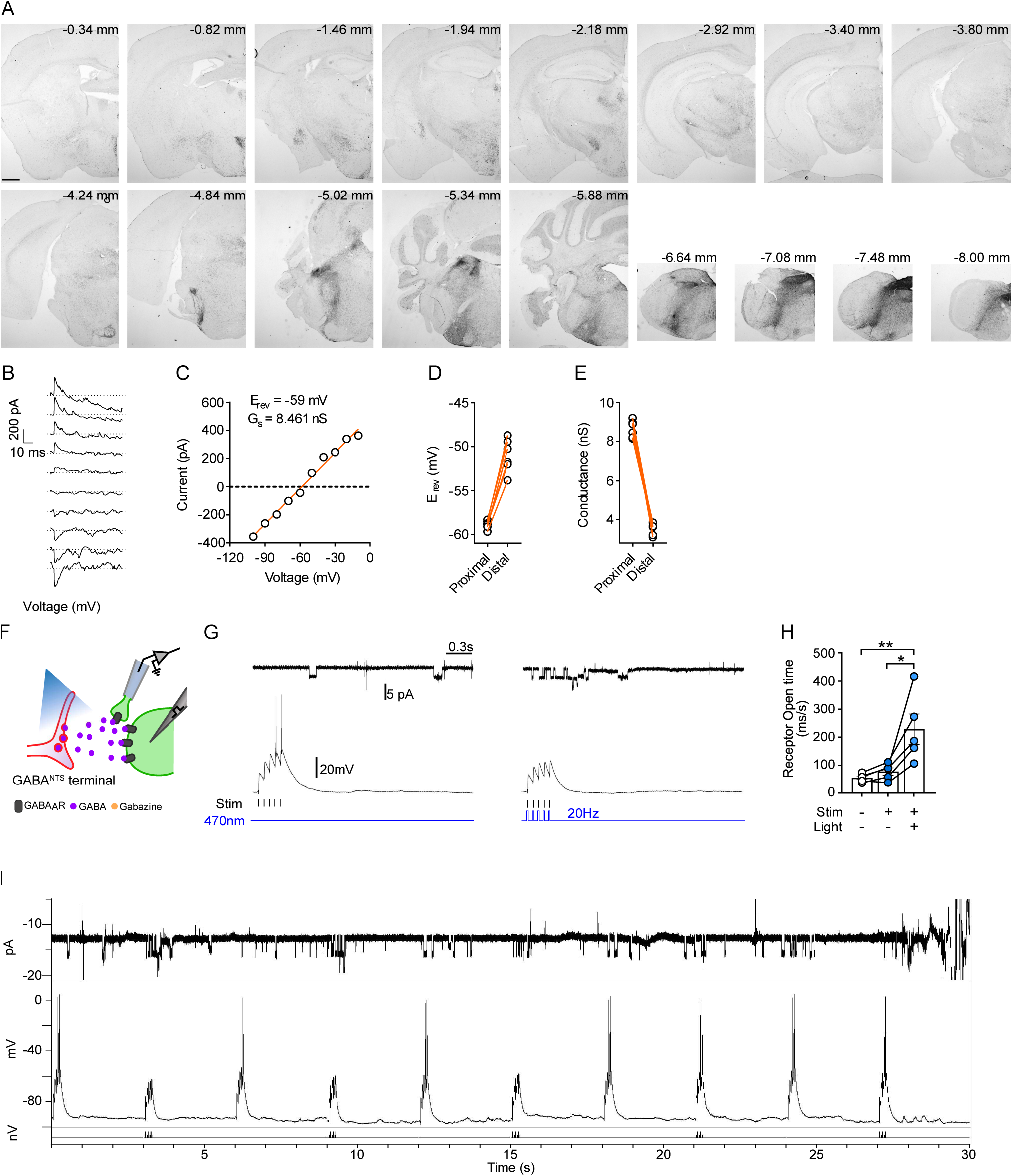
GABA^NTS^ projection pattern and CRACM additional analysis. (A) Representative photomicrographs (scale 100µm) of ChR2-mCherry fibers in a serial rostro-caudal distribution of a *Vgat^Cre^* mouse brain injected with AAV-DIO-ChR2-mCherry into the NTS. (B-E) A simulated voltage-clamp experiment in an anatomically realistic model of a medium spiny neuron, using the same voltage steps as those described for electrophysiology experiments (Figure 2E-G). The effect of optogenetic stimulation was modelled as a single GABAergic event taking place at each step. (C) Current amplitude was plotted against voltage (as in Figure 2E-G), and a linear fit was used to estimate peak conductance *G_s_* and reversal potential *E_rev_*. (D-E) Simulated optogenetic activation of GABA inputs making synaptic contact only on dendrites close to the soma (“prox.”, proximal dendrites) was compared to the same type of activation of GABA synapses connected only to dendrites far from the soma (“distal” dendrites) in six different medium spiny neuron models. The reversal potential for GABA in the model was set to -60 mV, and this is close to what was measured at the soma when GABA synaptic inputs were located on dendrites close to the soma. However, when GABA synapses were activated only on dendrites distant from the soma *E_rev_*, was more positive than expected and conductance *G_s_* was attenuated. (F) Diagram of outside-out patch technique coupled to a postsynaptic electrical stimulator. (G, left) Representative response of Npy^hrGFP^ cell subjected to series of five electrical stimuli, each evoking excitatory post-synaptic potential (EPSP) of sub-threshold amplitude and (G, right) same cell with electrical stimuli coupled with 470nm light burst. (M) Quantification of receptor opening time (n=5, two-way RM ANOVA F_(2,8)_=11.28; p=0.0047, Bonferroni adjusted P=0.0072 NS vs SL and P=0.0157 S vs SL), L-M n=4-5 cells. (I) Representative response of a single Npy^hrGFP^ cell subjected to 10 series of five electrical stimuli, each evoking excitatory post-synaptic potential (EPSP) of sub-threshold amplitude coupled with 470nm light burst in *Vgat^Cre^::NPY^hrGFP^* mouse injected with AAV-DIO-ChR2-mCherry into the NTS. NS: No stimulation; S: Stimulation; SL: Stimulation+Light.

**Figure S3.**
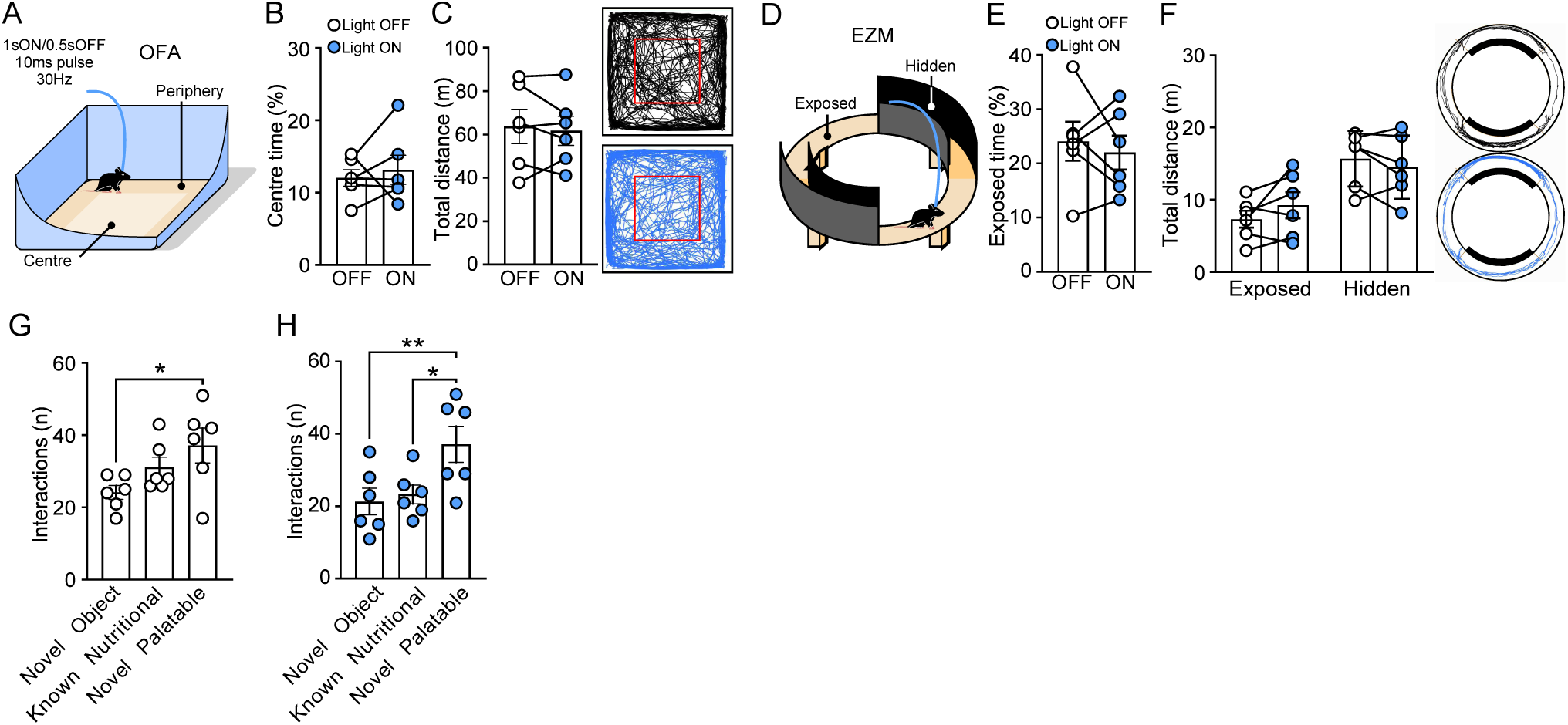
Optogenetic activation of GABA^NTS^→ARC in *Vgat^Cre^* mice does not induce anxiety-like behavior. (A) Diagram illustrating open field arena (OFA) task. GABA^NTS^→ARC does not alter (B) time spent in the center or (C) distance travelled (representative trace of the movement) during the test. (D) Diagram illustrating elevated zero maze (EZM) task. GABA^NTS^→ARC does not alter (E) time spent in the exposed area or (F) distance travelled in each zone (representative trace of the movement) during the test. (G-H) Quantification of number of interactions with a novel object, a known nutritional food item or a novel palatable food item in mice (G) non-stimulated (RM ANOVA (F_(2,10)_=4.993; p=0.0314, Bonferroni adjusted p=0.0306 NO vs NP) and (H) stimulated (RM ANOVA F_(2,10)_=10.71; p=0.0033, Bonferroni adjusted p=0.0051 NO vs NP; p=0.0121 KN vs NP). Data are expressed as individual values and as mean±S.E.M, n=6 mice. *:p<0.05; **:p<0.01. NO: novel object; NP: novel palatable; KN: known nutritional.

## REFERENCES

Aklan, I., Sayar Atasoy, N., Yavuz, Y., Ates, T., Coban, I., Koksalar, F., Filiz, G., Topcu, I. C., Oncul, M., Dilsiz, P., Cebecioglu, U., Alp, M. I., Yilmaz, B., Davis, D. R., Hajdukiewicz, K., Saito, K., Konopka, W., Cui, H., & Atasoy, D. (2020). NTS Catecholamine Neurons Mediate Hypoglycemic Hunger via Medial Hypothalamic Feeding Pathways. Cell Metabolism, 31(2), 313–326.e5. 10.1016/j.cmet.2019.11.016

Alexander, G. M., Rogan, S. C., Abbas, A. I., Armbruster, B. N., Pei, Y., Allen, J. A., Nonneman, R. J., Hartmann, J., Moy, S. S., Nicolelis, M. A., McNamara, J. O., & Roth, B. L. (2009). Remote control of neuronal activity in transgenic mice expressing evolved G protein-coupled receptors. Neuron, 63(1), 27–39. 10.1016/J.NEURON.2009.06.014

Alhadeff, A. L., Mergler, B. D., Zimmer, D. J., Turner, C. A., Reiner, D. J., Schmidt, H. D., Grill, H. J., & Hayes, M. R. (2017). Endogenous Glucagon-like Peptide-1 Receptor Signaling in the Nucleus Tractus Solitarius is Required for Food Intake Control. Neuropsychopharmacology : Official Publication of the American College of Neuropsychopharmacology, 42(7), 1471–1479. 10.1038/NPP.2016.246

Andermann, M. L., & Lowell, B. B. (2017). Toward a Wiring Diagram Understanding of Appetite Control. Neuron, 95(4), 757–778. 10.1016/J.NEURON.2017.06.014

Aponte, Y., Atasoy, D., & Sternson, S. M. (2011). AGRP neurons are sufficient to orchestrate feeding behavior rapidly and without training. Nature Neuroscience, 14(3), 351–355. 10.1038/NN.2739

Berridge, K. C. (2004). Motivation concepts in behavioral neuroscience. Physiology and Behavior, 81(2), 179–209. 10.1016/j.physbeh.2004.02.004

Berrios, J., Li, C., Madara, J. C., Garfield, A. S., Steger, J. S., Krashes, M. J., & Lowell, B. B. (2021). Food cue regulation of AGRP hunger neurons guides learning. Nature, 595, 695. 10.1038/s41586-021-03729-3

Betley, J. N., Cao, Z. F. H., Ritola, K. D., & Sternson, S. M. (2013). Parallel, Redundant Circuit Organization for Homeostatic Control of Feeding Behavior. Cell, 155(6), 1337–1350. 10.1016/J.CELL.2013.11.002

Betley, J. N., Xu, S., Cao, Z. F. H., Gong, R., Magnus, C. J., Yu, Y., & Sternson, S. M. (2015). Neurons for hunger and thirst transmit a negative-valence teaching signal. Nature, 521(7551), 180. 10.1038/NATURE14416

Blevins, J. E., Schwartz, M. W., & Baskin, D. G. (2004). Evidence that paraventricular nucleus oxytocin neurons link hypothalamic leptin action to caudal brain stem nuclei controlling meal size. American Journal of Physiology - Regulatory Integrative and Comparative Physiology, 287(1 56-1), 87–96. 10.1152/AJPREGU.00604.2003/ASSET/IMAGES/LARGE/ZH600704225 10005.JPEG

Campos, A., Port, J. D., & Acosta, A. (2022). Integrative Hedonic and Homeostatic Food Intake Regulation by the Central Nervous System: Insights from Neuroimaging. In Brain Sciences (Vol. 12, Issue 4). MDPI. 10.3390/brainsci12040431

Chen, H. Y., Trumbauer, M. E., Chen, A. S., Weingarth, D. T., Adams, J. R., Frazier, E. G., Shen, Z., Marsh, D. J., Feighner, S. D., Guan, X. M., Ye, Z., Nargund, R. P., Smith, R. G., van der Ploeg, L. H. T., Howard, A. D., Macneil, D. J., & Qian, S. (2004). Orexigenic Action of Peripheral Ghrelin Is Mediated by Neuropeptide Y and Agouti-Related Protein. Endocrinology, 145(6), 2607–2612. 10.1210/EN.2003-1596

Chen, J., Cheng, M., Wang, L., Zhang, L., Xu, D., Cao, P., Wang, F., Herzog, H., Song, S., & Zhan, C. (2020). A Vagal-NTS Neural Pathway that Stimulates Feeding. Current Biology, 30(20), 3986–3998.e5. 10.1016/J.CUB.2020.07.084

Cheng, W., Gordian, D., Ludwig, M. Q., Pers, T. H., Seeley, R. J., & Myers, M. G. (2022). Hindbrain circuits in the control of eating behaviour and energy balance. Nature Metabolism, 4(7), 826–835. 10.1038/s42255-022-00606-9

Cheng, W., Ndoka, E., Hutch, C., Roelofs, K., MacKinnon, A., Khoury, B., Magrisso, J., Kim, K. S., Rhodes, C. J., Olson, D. P., Seeley, R. J., Sandoval, D., & Myers, M. G. (2020). Leptin receptor-expressing nucleus tractus solitarius neurons suppress food intake independently of GLP1 in mice. JCI Insight, 5(7). 10.1172/JCI.INSIGHT.134359

Clifton, P. G. (2000). Meal patterning in rodents: psychopharmacological and neuroanatomical studies. Neuroscience & Biobehavioral Reviews, 24(2), 213–222. 10.1016/S0149-7634(99)00074-3

Cork, S. C., Richards, J. E., Holt, M. K., Gribble, F. M., Reimann, F., & Trapp, S. (2015). Distribution and characterisation of Glucagon-like peptide-1 receptor expressing cells in the mouse brain. Molecular Metabolism, 4(10), 718–731. 10.1016/j.molmet.2015.07.008

Cummings, D. E., Purnell, J. Q., Frayo, R. S., Schmidova, K., Wisse, B. E., & Weigle, D. S. (2001). A Preprandial Rise in Plasma Ghrelin Levels Suggests a Role in Meal Initiation in Humans. Diabetes, 50(8), 1714–1719. 10.2337/DIABETES.50.8.1714

D’Agostino, G., & Luckman, S. M. (2022). Brainstem peptides and peptidergic neurons in the regulation of appetite. Current Opinion in Endocrine and Metabolic Research, 24, 100339. 10.1016/J.COEMR.2022.100339

D’Agostino, G., Lyons, D., Cristiano, C., Lettieri, M., Olarte-Sanchez, C., Burke, L. K., Greenwald-Yarnell, M., Cansell, C., Doslikova, B., Georgescu, T., Martinez de Morentin, P. B., Myers, M. G., Rochford, J. J., & Heisler, L. K. (2018). Nucleus of the Solitary Tract Serotonin 5-HT2C Receptors Modulate Food Intake. Cell Metabolism, 28(4), 619–630.e5. 10.1016/J.CMET.2018.07.017

D’Agostino, G., Lyons, D. J., Cristiano, C., Burke, L. K., Madara, J. C., Campbell, J. N., Garcia, A. P., Land, B. B., Lowell, B. B., Dileone, R. J., & Heisler, L. K. (2016). Appetite controlled by a cholecystokinin nucleus of the solitary tract to hypothalamus neurocircuit. ELife, 5. 10.7554/eLife.12225

Deem, J. D., Faber, C. L., & Morton, G. J. (2022). AgRP neurons: Regulators of feeding, energy expenditure, and behavior. FEBS Journal, 289(8), 2362–2381. 10.1111/FEBS.16176

Dietrich, M. O., Bober, J., Ferreira, J. G., Tellez, L. A., Mineur, Y. S., Souza, D. O., Gao, X.- B., Picciotto, M. R., Araújo, I., Liu, Z.-W., & Horvath, T. L. (2012). AgRP neurons regulate development of dopamine neuronal plasticity and nonfood-associated behaviors. Nature Neuroscience. 10.1038/nn.3147

Dietrich, M. O., Zimmer, M. R., Bober, J., & Horvath, T. L. (2015). Hypothalamic Agrp Neurons Drive Stereotypic Behaviors beyond Feeding. Cell, 160(6), 1222–1232. 10.1016/J.CELL.2015.02.024

Dowsett, G. K. C., Lam, B. Y. H., Tadross, J. A., Cimino, I., Rimmington, D., Coll, A. P., Polex-Wolf, J., Knudsen, L. B., Pyke, C., & Yeo, G. S. H. (2021). A survey of the mouse hindbrain in the fed and fasted states using single-nucleus RNA sequencing. Molecular Metabolism, 53, 101240. 10.1016/J.MOLMET.2021.101240

Fenno, L. E., Mattis, J., Ramakrishnan, C., & Deisseroth, K. (2017). A Guide to Creating and Testing New INTRSECT Constructs. Current Protocols in Neuroscience, 80(1), 4.39.1–4.39.24. 10.1002/CPNS.30

Fenno, L. E., Mattis, J., Ramakrishnan, C., Hyun, M., Lee, S. Y., He, M., Tucciarone, J., Selimbeyoglu, A., Berndt, A., Grosenick, L., Zalocusky, K. A., Bernstein, H., Swanson, H., Perry, C., Diester, I., Boyce, F. M., Bass, C. E., Neve, R., Huang, Z. J., & Deisseroth, K. (2014). Targeting cells with single vectors using multiple-feature Boolean logic. Nature Methods, 11(7), 763–772. 10.1038/nmeth.2996

Fortin, S. M., Lipsky, R. K., Lhamo, R., Chen, J., Kim, E., Borner, T., Schmidt, H. D., & Hayes, M. R. (2020). GABA neurons in the nucleus tractus solitarius express GLP-1 receptors and mediate anorectic effects of liraglutide in rats HHS Public Access. Sci Transl Med, 12(533), 109–114. 10.1126/scitranslmed.aay8071

Garau, C., Blomeley, C., Burdakov, D., & Burdakov, D. (2020). Orexin neurons and inhibitory Agrp→orexin circuits guide spatial exploration in mice. The Journal of Physiology, 598, 4371–4383. 10.1113/JP280158

Georgescu, T., Lyons, D., Doslikova, B., Garcia, A. P., Marston, O., Burke, L. K., Chianese, R., Lam, B. Y. H., Yeo, G. S. H., Rochford, J. J., Garfield, A. S., & Heisler, L. K. (2020). Neurochemical Characterization of Brainstem Pro-Opiomelanocortin Cells. Endocrinology, 161(4). 10.1210/endocr/bqaa032

González, J. A., Iordanidou, P., Strom, M., Adamantidis, A., & Burdakov, D. (2016). Awake dynamics and brain-wide direct inputs of hypothalamic MCH and orexin networks. Nature Communications, 7. 10.1038/NCOMMS11395

Graham, D. L., Durai, H. H., Trammell, T. S., Noble, B. L., Mortlock, D. P., Galli, A., & Stanwood, G. D. (2020). A novel mouse model of glucagon-like peptide-1 receptor expression: A look at the brain. Journal of Comparative Neurology, 528(14), 2445–2470. 10.1002/cne.24905

Grill, H. J., & Hayes, M. R. (2012). Hindbrain Neurons as an Essential Hub in the Neuroanatomically Distributed Control of Energy Balance. Cell Metabolism, 16(3), 296–309. 10.1016/J.CMET.2012.06.015

Hahn, T. M., Breininger, J. F., Baskin, D. G., & Schwartz, M. W. (1998). Coexpression of Agrp and NPY in fasting-activated hypothalamic neurons. Nature Neuroscience 1998 1:4, 1(4), 271–272. 10.1038/1082

Han, Y., Xia, G., Srisai, D., Meng, F., He, Y., Ran, Y., He, Y., Farias, M., Hoang, G., Tóth, I., Dietrich, M. O., Chen, M. H., Xu, Y., & Wu, Q. (2021). Deciphering an AgRP-serotoninergic neural circuit in distinct control of energy metabolism from feeding. Nature Communications 2021 12:1, 12(1), 1–16. 10.1038/s41467-021-23846-x

Hao, Y., Hao, S., Andersen-Nissen, E., Mauck, W. M., Zheng, S., Butler, A., Lee, M. J., Wilk, A. J., Darby, C., Zager, M., Hoffman, P., Stoeckius, M., Papalexi, E., Mimitou, E. P., Jain, J., Srivastava, A., Stuart, T., Fleming, L. M., Yeung, B., … Satija, R. (2021). Integrated analysis of multimodal single-cell data. Cell, 184(13), 3573–3587.e29. 10.1016/j.cell.2021.04.048

Heinz, D. E., Schöttle, V. A., Nemcova, P., Binder, F. P., Ebert, T., Domschke, K., & Wotjak, C. T. (2021). Exploratory drive, fear, and anxiety are dissociable and independent components in foraging mice. Translational Psychiatry 2021 11:1, 11(1), 1–12. 10.1038/s41398-021-01458-9

Heisler, L. K., Jobst, E. E., Sutton, G. M., Zhou, L., Borok, E., Thornton-Jones, Z., Liu, H. Y., Zigman, J. M., Balthasar, N., Kishi, T., Lee, C. E., Aschkenasi, C. J., Zhang, C. Y., Yu, J., Boss, O., Mountjoy, K. G., Clifton, P. G., Lowell, B. B., Friedman, J. M., … Cowley, M. A. (2006). Serotonin Reciprocally Regulates Melanocortin Neurons to Modulate Food Intake. Neuron, 51(2), 239–249. 10.1016/j.neuron.2006.06.004

Heisler, L. K., & Lam, D. D. (2017). An appetite for life: brain regulation of hunger and satiety. Current Opinion in Pharmacology, 37, 100–106. 10.1016/J.COPH.2017.09.002

Hentges, S. T., Otero-Corchon, V., Pennock, R. L., King, C. M., & Low, M. J. (2009). Proopiomelanocortin Expression in both GABA and Glutamate Neurons. The Journal of Neuroscience, 29(43), 13684–13690. 10.1523/JNEUROSCI.3770-09.2009

Holt, M. K., Richards, J. E., Cook, D. R., Brierley, D. I., Williams, D. L., Reimann, F., Gribble, F. M., & Trapp, S. (2019). Preproglucagon neurons in the nucleus of the solitary tract are the main source of brain GLP-1, mediate stress-induced hypophagia, and limit unusually large intakes of food. Diabetes, 68(1), 21–33. 10.2337/DB18-0729/-/DC1

Hyun, U., & Sohn, J. W. (2022). Autonomic control of energy balance and glucose homeostasis. Experimental & Molecular Medicine 2022 54:4, 54(4), 370–376. 10.1038/s12276-021-00705-9

Jensen, C. B., Pyke, C., Rasch, M. G., Dahl, A. B., Knudsen, L. B., & Secher, A. (2018). Characterization of the Glucagonlike Peptide-1 Receptor in Male Mouse Brain Using a Novel Antibody and In Situ Hybridization. Endocrinology, 159(2), 665–675. 10.1210/en.2017-00812

Jikomes, N., Ramesh, R. N., Mandelblat-Cerf, Y., & Andermann, M. L. (2016). Preemptive Stimulation of AgRP Neurons in Fed Mice Enables Conditioned Food Seeking under Threat. Current Biology, 26(18), 2500–2507. 10.1016/J.CUB.2016.07.019

Kim, S. Y., Adhikari, A., Lee, S. Y., Marshel, J. H., Kim, C. K., Mallory, C. S., Lo, M., Pak, S., Mattis, J., Lim, B. K., Malenka, R. C., Warden, M. R., Neve, R., Tye, K. M., & Deisseroth, K. (2013). Diverging neural pathways assemble a behavioural state from separable features in anxiety. Nature 2013 496:7444, 496(7444), 219–223. 10.1038/nature12018

Krashes, M. J., Koda, S., Ye, C. P., Rogan, S. C., Adams, A. C., Cusher, D. S., Maratos-Flier, E., Roth, B. L., & Lowell, B. B. (2011). Rapid, reversible activation of AgRP neurons drives feeding behavior in mice. Journal of Clinical Investigation, 121(4), 1424–1428. 10.1172/JCI46229

Krashes, M. J., Shah, B. P., Madara, J. C., Olson, D. P., Strochlic, D. E., Garfield, A. S., Vong, L., Pei, H., Watabe-Uchida, M., Uchida, N., Liberles, S. D., & Lowell, B. B. (2014). An excitatory paraventricular nucleus to AgRP neuron circuit that drives hunger. Nature 2014 507:7491, 507(7491), 238–242. 10.1038/nature12956

Krnjević, K., & Schwartz, S. (1967). The action of γ-Aminobutyric acid on cortical neurones. Experimental Brain Research, 3(4), 320–336. 10.1007/BF00237558

Li, C., Hou, Y., Zhang, J., Sui, G., Du, X., Licinio, J., Wong, M. L., & Yang, Y. (2019). AGRP neurons modulate fasting-induced anxiolytic effects. Translational Psychiatry 2019 9:1, 9(1), 1–10. 10.1038/s41398-019-0438-1

Lin, J. Y., Lin, M. Z., Steinbach, P., & Tsien, R. Y. (2009). Characterization of Engineered Channelrhodopsin Variants with Improved Properties and Kinetics. Biophysical Journal, 96(5), 1803–1814. 10.1016/J.BPJ.2008.11.034

Lindroos, R., & Hellgren Kotaleski, J. (2021). Predicting complex spikes in striatal projection neurons of the direct pathway following neuromodulation by acetylcholine and dopamine. The European Journal of Neuroscience, 53(7), 2117–2134. 10.1111/ejn.14891

Liu, J., Conde, K., Zhang, P., Lilascharoen, V., Xu, Z., Lim, B. K., Seeley, R. J., Zhu, J. J., Scott, M. M., & Pang, Z. P. (2017). Enhanced AMPA receptor trafficking mediates the anorexigenic effect of endogenous glucagon like peptide-1 in the paraventricular hypothalamus. Neuron, 96(4), 897. 10.1016/J.NEURON.2017.09.042

Ludwig, M. Q., Cheng, W., Gordian, D., Lee, J., Paulsen, S. J., Hansen, S. N., Egerod, K. L., Barkholt, P., Rhodes, C. J., Secher, A., Knudsen, L. B., Pyke, C., Myers, M. G., & Pers, T. H. (2021). A genetic map of the mouse dorsal vagal complex and its role in obesity. Nature Metabolism, 3(4), 530–545. 10.1038/s42255-021-00363-1

Luquet, S., Perez, F. A., Hnasko, T. S., & Palmiter, R. D. (2005). NPY/AgRP neurons are essentials for feeding in adult mice but can be ablated in neonates. Science, 310(5748), 683–685. 10.1126/SCIENCE.1115524

Luquet, S., Phillips, C. T., & Palmiter, R. D. (2007). NPY/AgRP neurons are not essential for feeding responses to glucoprivation. Peptides, 28(2), 214–225. 10.1016/J.PEPTIDES.2006.08.036

Madisen, L., Zwingman, T. A., Sunkin, S. M., Oh, S. W., Zariwala, H. A., Gu, H., Ng, L. L., Palmiter, R. D., Hawrylycz, M. J., Jones, A. R., Lein, E. S., & Zeng, H. (2009). A robust and high-throughput Cre reporting and characterization system for the whole mouse brain. Nature Neuroscience 2009 13:1, 13(1), 133–140. 10.1038/nn.2467

Mandelblat-Cerf, Y., Ramesh, R. N., Burgess, C. R., Patella, P., Yang, Z., Lowell, B. B., & Andermann, M. L. (2015). Arcuate hypothalamic AgRP and putative POMC neurons show opposite changes in spiking across multiple timescales. ELife, 4. 10.7554/eLife.07122

Olson, B. R., Freilino, M., Hoffman, G. E., Stricker, E. M., Sved, A. F., & Verbalis, J. G. (1993). C-Fos expression in rat brain and brainstem nuclei in response to treatments that alter food intake and gastric motility. Molecular and Cellular Neurosciences, 4(1), 93–106. 10.1006/MCNE.1993.1011

Petreanu, L., Huber, D., Sobczyk, A., & Svoboda, K. (2007). Channelrhodopsin-2-assisted circuit mapping of long-range callosal projections. 10.1038/nn1891

Richard, C. D., Tolle, V., & Low, M. J. (2011). Meal pattern analysis in neural-specific proopiomelanocortin-deficient mice. European Journal of Pharmacology, 660(1), 131–138. 10.1016/J.EJPHAR.2010.12.022

Shi, M. Y., Ding, L. F., Guo, Y. H., Cheng, Y. X., Bi, G. Q., & Lau, P. M. (2021). Long-range GABAergic projections from the nucleus of the solitary tract. Molecular Brain, 14(1), 1–5. 10.1186/S13041-021-00751-4/FIGURES/1

Spruston, N., Jaffe, D. B., Williams, S. H., & Johnston, D. (1993). Voltage- and space-clamp errors associated with the measurement of electrotonically remote synaptic events. Journal of Neurophysiology, 70(2), 781–802. 10.1152/jn.1993.70.2.781

Stamatakis, A. M., & Stuber, G. D. (2012). Activation of lateral habenula inputs to the ventral midbrain promotes behavioral avoidance. Nature Neuroscience 2012 15:8, 15(8), 1105–1107. 10.1038/nn.3145

Sylantyev, S., Savtchenko, L. P., O’Neill, N., & Rusakov, D. A. (2020). Extracellular GABA waves regulate coincidence detection in excitatory circuits. The Journal of Physiology, 598(18), 4047. 10.1113/JP279744

Tsang, A. H., Nuzzaci, D., Darwish, T., Samudrala, H., & Blouet, C. (2020). Nutrient sensing in the nucleus of the solitary tract mediates non-aversive suppression of feeding via inhibition of AgRP neurons. Molecular Metabolism, 42, 101070. 10.1016/J.MOLMET.2020.101070

van den Pol, A. N., Yao, Y., Fu, L. Y., Foo, K., Huang, H., Coppari, R., Lowell, B. B., & Broberger, C. (2009). Neuromedin B and gastrin-releasing peptide excite arcuate nucleus neuropeptide Y neurons in a novel transgenic mouse expressing strong Renilla green fluorescent protein in NPY neurons. The Journal of Neuroscience : The Official Journal of the Society for Neuroscience, 29(14), 4622–4639. 10.1523/JNEUROSCI.3249-08.2009

Vong, L., Ye, C., Yang, Z., Choi, B., Chua, S., & Lowell, B. B. (2011). Leptin action on GABAergic neurons prevents obesity and reduces inhibitory tone to POMC neurons. Neuron, 71(1), 142–154. 10.1016/J.NEURON.2011.05.028

Wagner, S., Brierley, D. I., Leeson-Payne, A., Jiang, W., Chianese, R., Lam, B. Y. H., Dowsett, G. K. C., Cristiano, C., Lyons, D., Reimann, F., Gribble, F. M., Martinez de Morentin, P. B., Yeo, G. S. H., Trapp, S., & Heisler, L. K. (2022). Obesity medication lorcaserin activates brainstem GLP-1 neurons to reduce food intake and augments GLP-1 receptor agonist induced appetite suppression. Molecular Metabolism, 68. 10.1016/J.MOLMET.2022.101665

Wang, C., Zhou, W., He, Y., Yang, T., Xu, P., Yang, Y., Cai, X., Wang, J., Liu, H., Yu, M., Liang, C., Yang, T., Liu, H., Fukuda, M., Tong, Q., Wu, Q., Sun, Z., He, Y., & Xu, Y. (2021). AgRP neurons trigger long-term potentiation and facilitate food seeking. Translational Psychiatry, 11, 11. 10.1038/s41398-020-01161-1

Williams, D. L., & Schwartz, M. W. (2005). The melanocortin system as a central integrator of direct and indirect controls of food intake. American Journal of Physiology. Regulatory, Integrative and Comparative Physiology, 289(1). 10.1152/AJPREGU.00226.2005

Wu, Q., Lemus, M. B., Stark, R., Bayliss, J. A., Reichenbach, A., Lockie, S. H., & Andrews, Z. B. (2014). The Temporal Pattern of cfos Activation in Hypothalamic, Cortical, and Brainstem Nuclei in Response to Fasting and Refeeding in Male Mice. Endocrinology, 155(3), 840–853. 10.1210/EN.2013-1831

Yavari, A., Stocker, C. J., Ghaffari, S., Wargent, E. T., Steeples, V., Czibik, G., Pinter, K., Bellahcene, M., Woods, A., Martinez de Morentin, P. B., Cansell, C., Lam, B. Y., Chuster, A., Petkevicius, K., Nguyen-Tu, M. S., Martinez-Sanchez, A., Pullen, T. J., Oliver, P. L., Stockenhuber, A., … Ashrafian, H. (2016). Chronic Activation of gamma2 AMPK Induces Obesity and Reduces beta Cell Function. Cell Metab, 23(5), 821–836. 10.1016/j.cmet.2016.04.003

Zhan, C., Zhou, J., Feng, Q., Zhang, J. en, Lin, S., Bao, J., Wu, P., & Luo, M. (2013). Acute and long-term suppression of feeding behavior by POMC neurons in the brainstem and hypothalamus, respectively. The Journal of Neuroscience : The Official Journal of the Society for Neuroscience, 33(8), 3624–3632. 10.1523/JNEUROSCI.2742-12.2013

Zingg, B., Chou, X. lin, Zhang, Z. gang, Mesik, L., Liang, F., Tao, H. W., & Zhang, L. I. (2017). AAV-Mediated Anterograde Transsynaptic Tagging: Mapping Corticocollicular Input-Defined Neural Pathways for Defense Behaviors. Neuron, 93(1), 33–47. 10.1016/J.NEURON.2016.11.045

Zorrilla, E. P., Inoue, K., Fekete, É. M., Tabarin, A., Valdez, G. R., & Koob, G. F. (2005). Measuring meals: Structure of prandial food and water intake of rats. American Journal of Physiology - Regulatory Integrative and Comparative Physiology, *288*(6 57-6), 1450–1467. 10.1152/AJPREGU.00175.2004/SUPPL_FILE/ZORRILLA_SUPP_TABLE.PDF

